# Antibiotic killing of drug-induced bacteriostatic cells

**DOI:** 10.1101/2024.09.06.611640

**Authors:** Teresa Gil-Gil, Brandon A. Berryhill

## Abstract

**Background:** There is a long-standing belief that bacteriostatic drugs are inherently antagonistic to the action of bactericidal antibiotics. This belief is primarily due to the fact that the action of most bactericidal antibiotics requires the target bacteria to be growing. Since bacteriostatic drugs stop the growth of treated bacteria, these drugs would necessarily work against one another. We have recently shown that bacteria treated with high concentrations of bacteriostatic drugs retain some metabolic activity, dividing on average once per day.

**Objectives:** We seek to determine if this low level of growth is sufficient to allow for bactericidal antibiotics of different classes to still kill after bacteria are treated with bacteriostatic drugs.

**Methods:** We first treated *Escherichia coli* and *Staphylococcus aureus* with two different bacteriostatic drugs, followed by one of three bactericidal drugs of three different classes. The density of these bacteria was tracked over six days to determine the amount of killing that occurred.

**Results:** Our results question this long-standing belief by demonstrating conditions where sequential treatment with a bacteriostatic then bactericidal antibiotic is as or more effective than treatment with a bactericidal drug alone.

**Conclusions:** These results raise the need to investigate the pharmacodynamics of the joint action of bacteriostatic and bactericidal antibiotics *in vitro* and *in vivo*.

## Introduction

The clinical outcomes of antibiotic treatments involving the administration of bacteriostatic (which inhibit bacterial growth) or bactericidal (which kill bacteria) antibiotics are complex and context-dependent ^1,2^. While antibiotic co-administration strategies can be beneficial in specific contexts, they require careful consideration regarding the interaction of the drugs, individualized patient factors, infection types, and resistance patterns ^3^. Therefore, it is necessary to evaluate specific drug combinations rather than relying solely on their classification as bacteriostatic or bactericidal. Historically, the co-administration of bacteriostatic and bactericidal antibiotics has been discouraged due to the notion that bactericidal drugs require actively dividing, metabolically active bacteria to exert their effects—an action that is directly opposed by bacteriostatic drugs ^4^. Despite this anticipated antagonism, many multi-drug treatment regimens employed for difficult-to-treat infections use combinations of both of these drugs, often to some degree of success ^5^. International guidelines often recommend combining bacteriostatic drugs, such as third-generation tetracyclines (e.g., tigecycline) or oxazolidinones (e.g., linezolid), with bactericidal agents as a last-resort treatment option ^6,7^. Thus, these antibiotics cannot be purely antagonistic as commonly believed. Recent work by Gil-Gil et al. has shown that, even under high concentrations of bacteriostatic drugs, cell division occurs albeit at a rate nearly 100 times slower than in untreated cultures ^8^. Taken together, these previously reported results provide potential evidence and a mechanism that runs counter to the maxim that bacteriostatic and bactericidal antibiotics should not be used together.

Here, we use a laboratory strain of *Escherichia coli* ^9^ and a clinical isolate of *Staphylococcus aureus* ^10^ with combinations of bacteriostatic and bactericidal drugs to determine how pre-exposure to bacteriostatic drugs changes the dynamics of subsequent treatment with bactericidal agents. Surprisingly, we did not observe any conditions that resulted in complete antagonism between bacteriostatic and bactericidal drugs. Instead, our findings revealed a spectrum of outcomes, including delayed culture clearance, accelerated culture clearance, and even prevention of resistance emergence. These results question the maxim that bacteriostatic and bactericidal antibiotics should not be used together and raise the need to further investigate the pharmacodynamics of the use of both of these drug types together.

## Results

### Determining the concentration of bacteriostatic antibiotics

To open our exploration of the dynamics of joint treatment with bacteriostatic and bactericidal antibiotics, we first determined the Minimum Inhibitory Concentration (MIC) of each drug to the two bacteria used for this study, *E. coli* MG1655 and *S. aureus* MN8. The MIC values for *E. coli* were 6.25 μg/mL for chloramphenicol, 25 μg/mL for azithromycin, 0.03 μg/mL for ciprofloxacin, 12 μg/mL for gentamicin, and 25 μg/mL for ampicillin. For *S. aureus*, the MIC values were 8 μg/mL for chloramphenicol, 0.15 μg/mL for tetracycline, 0.15 μg/mL for ciprofloxacin, 1 μg/mL for gentamicin, and 256 μg/mL for ampicillin. We then determined the concentration of the bacteriostatic antibiotic that prevents net growth over the course of the 6-day experiment while not causing a substantial amount of death in these cultures (Figure 1). Based on these results, all further experiments were conducted using 4x MIC concentrations of chloramphenicol and azithromycin for *E. coli* and 4x MIC for chloramphenicol and 10x MIC for tetracycline when using *S. aureus*.

**Figure 1.**
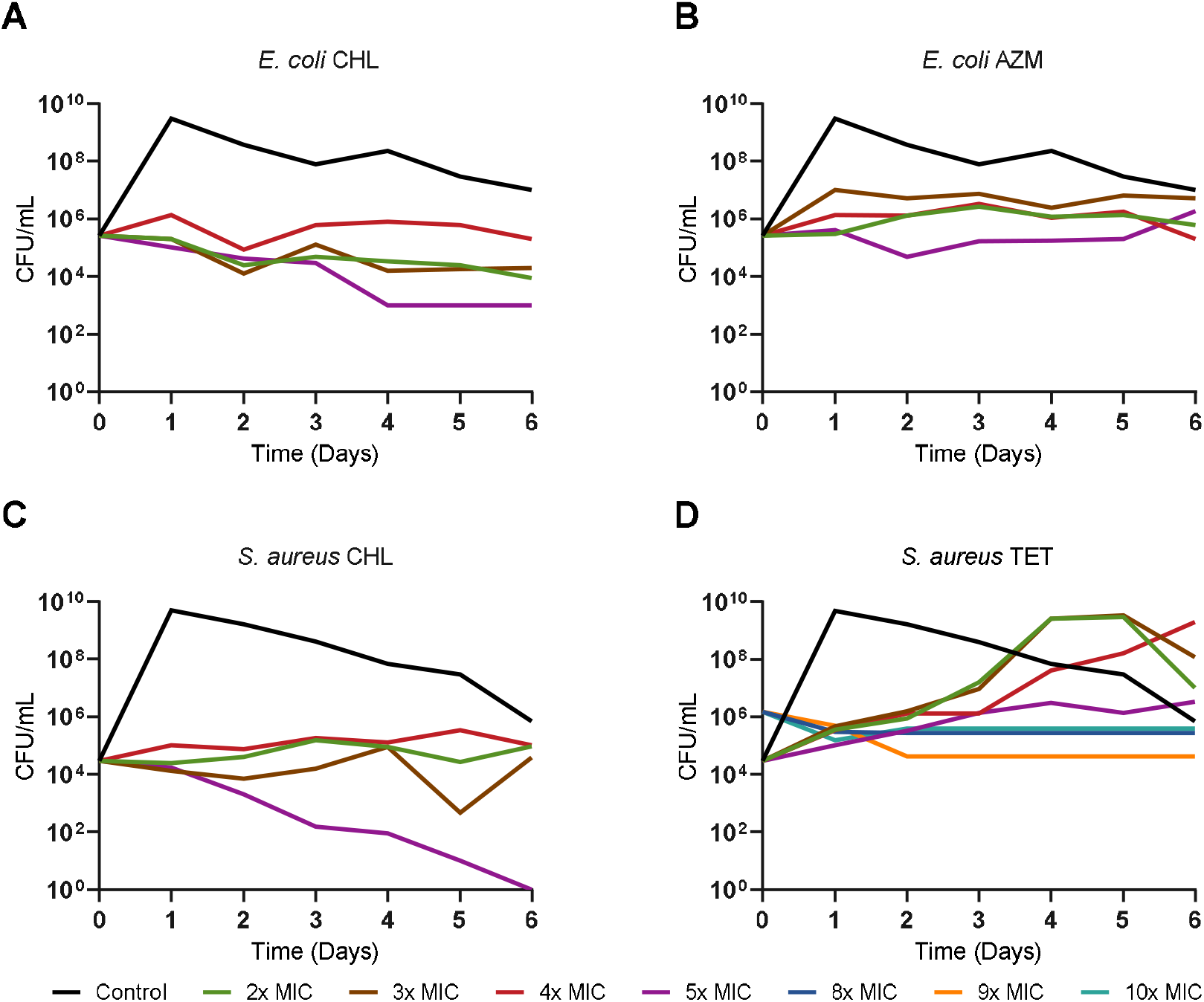
Exposure to varying concentrations of bacteriostatic drugs for six days without transferring. Density in CFU/mL of *E. coli* MG1655 (A and B) and *S. aureus* MN8 (C and D) measured every day for six days at several concentrations of either chloramphenicol (CHL) (A and C), azithromycin (AZM) (B), or tetracycline (TET) (D). Shown in black for each panel is a control culture without antibiotics.

### Treatment with a bacteriostatic then a bactericidal antibiotic

We initiated our joint action experiments by treating *E. coli* MG1655 with either chloramphenicol or azithromycin at a super-MIC concentration for one day. Following pre-treatment, we introduced one of several bactericidal drugs with differing mechanisms of action (Figure 2A, B, and C). Cultures were sampled each daily over a 6-day period. In every scenario where both drugs were used sequentially, the bacterial density at day 6 was consistently lower than in cultures containing just the bacteriostatic drug, indicating that bactericidal activity occurred. Notably, when gentamicin was administered after the bacteriostatic pre-treatment, bacterial clearance occurred faster than when gentamicin was used alone. This accelerated clearance is likely because, without the bacteriostatic pre-treatment, small colony variants resistant to gentamicin emerge; the pre-treatment effectively prevented the appearance of these resistant variants – thereby, enhancing the overall efficacy of gentamicin. However, the dynamics of treatment with ciprofloxacin and ampicillin following the bacteriostatic drug were slower than those when either ciprofloxacin or ampicillin were used alone.

**Figure 2.**
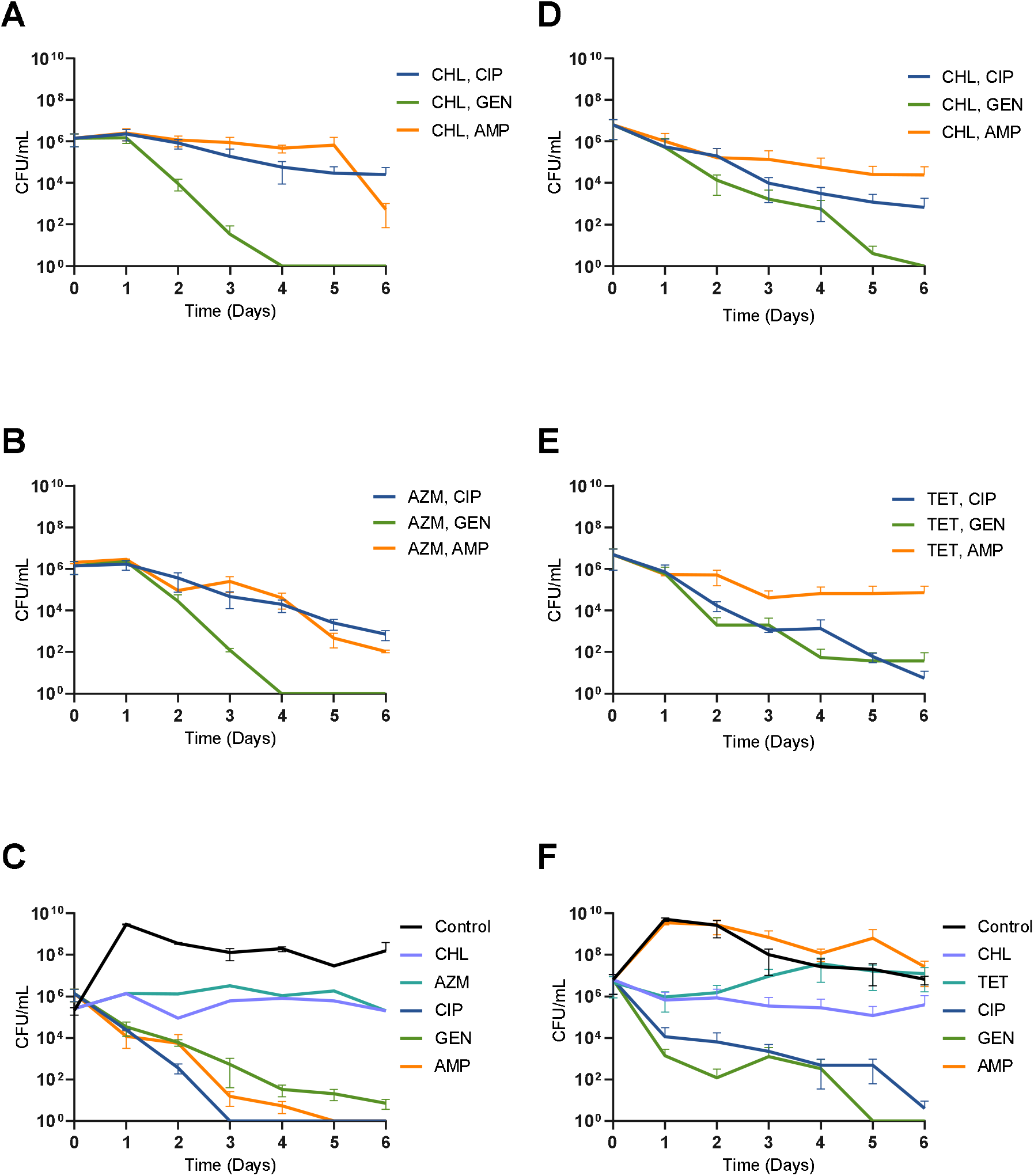
The treatment of *E. coli* and *S. aureus* with bacteriostatic followed by bactericidal antibiotics. Shown are bacterial densities in CFU/mL, each point represents the mean and standard deviation of three biological replicates. Cultures are sampled every day for six days and are not transferred. A, B, and C correspond to *E. coli* experiments and D, E, and F to *S. aureus* experiments. Chloramphenicol is at 4x MIC, azithromycin at 4x MIC, tetracycline is at 10x MIC, and the bactericidal drugs are all at 4x MIC for *E. coli* and 12x MIC for *S. aureus*. For panels A, B, D, and E cultures were treated with either chloramphenicol, azithromycin, or tetracycline for one day before the subsequent addition of a bactericidal antibiotic. Figure C and F are a control where cultures were treated with only a bactericidal or bacteriostatic drug.

When a clinical isolate of *S. aureus* was treated first with either chloramphenicol or tetracycline followed by a bactericidal drug, the results obtained are qualitatively like those obtained with *E. coli* (Figure 2C, D, and E). Interestingly, the combination of bacteriostatic drugs followed by ampicillin led to significantly greater culture clearance than ampicillin alone. While both bacteriostatic drugs combined with gentamicin or ciprofloxacin resulted in high levels of bacterial killing, these combinations slowed down the dynamics of both bactericidal drugs.

## Discussion

Contrary to the maxim that bacteriostatic drugs are completely antagonistic to bactericidal agents ^4^, our findings indicate that bactericidal action does occur after bacteriostatic pre-treatment. In neither *E. coli* nor *S. aureus*, pre-treatment with bacteriostatic antibiotics was completely antagonistic to the subsequent action of bactericidal drugs. Quite to the contrary, pre-treatment with bacteriostatic drugs in several cases allowed for a higher degree of killing than treatment with just the bactericidal agents alone. Two key observations emerged from our study. First, bacteriostatic pre-treatment effectively suppressed the emergence of gentamicin-resistant small colony variants in *E. coli* that ascend with the treatment of gentamicin alone, increasing the rate of bacterial clearance ^11^. Second, in the case of *S. aureus* MN8, an inducible beta-lactamase producer, pre-treatment with bacteriostatic drugs facilitated better control of bacterial density by ampicillin, although a significant population of what are likely persister cells remained. These results are consistent with the concept of cellular hysteresis and open the door to looking at collateral susceptibility with parings of bacteriostatic and bactericidal antibiotics ^12,13^.

As with all *in vitro* studies of antibiotics, the role of the host’s immune system was not considered in our experiments, a critical factor in the success of antibiotic therapy ^14^. One hypothesis regarding why antibiotic therapy is successful *in vivo* is that the drugs slow down growth or reduce the density of bacteria, providing an opportunity for the host’s innate immune response to control and potentially eliminate the infecting bacteria ^15^. While our study did not achieve complete bacterial clearance in all scenarios, whether through single-drug therapy or combined bacteriostatic and bactericidal treatments, we suggest that stopping the net growth of bacteria and achieving a higher degree of clearance over time would be sufficient for the host’s immune system to control the residual bacterial population effectively. Subsequently, it is essential to further explore the pharmacodynamics of combined bacteriostatic and bactericidal treatments, both *in vitro* and *in vivo*, to better design antibiotic treatment protocols.

## Materials and Methods

### Growth media

LB broth (244620) was obtained from BD. Muller Hinton II (MHII) Broth (90922-500G) obtained from Millipore. LB agar (244510) for plates was obtained from BD.

### Growth conditions

All experiments were conducted at 37°C and shaken continuously.

### Bacterial strains

*E. coli* MG1655 was obtained from the Levin Lab Bacterial collection. *Staphylococcus aureus* MN8 was obtained from Tim Read of Emory University.

### Antibiotics

Ciprofloxacin (A4556) was obtained from AppliChem Panreac, Chloramphenicol (23660) was obtained from USB, Ampicillin (A9518-25G) was obtained from Sigma Aldrich, Azithromycin (3771) was obtained from TOCRIS, Gentamicin (BP918-1) was obtained from Fisher BioReagents, and Tetracycline (T17999) was obtained from Research Products International.

### Sampling bacterial densities

The densities of bacteria were estimated by serial dilution in 0.85% saline and plating. The total density of bacteria was estimated on Lysogeny Broth (LB) plates with 1.6% agar.

### Minimum inhibitory concentrations

Antibiotic MICs were estimated using a 2-fold microdilution procedure as described in ^16^.

### Antibiotic killing assays

For these experiments, overnight cultures of *E. coli* MG1655 or *S. aureus* MN8 were added to LB broth or MHII at an initial density of approximately 10^6^ cells per mL, followed by a 24 h incubation with a bacteriostatic drug at the concentrations indicated in the figure captions. After the 24 h incubation, a bactericidal antibiotic was added at the concentrations indicated in the figure captions. The cultures containing both drugs were incubated for five days without transferring. Bacterial densities were estimated before the addition of the bacteriostatic drugs (t=0), before the addition of the bactericidal drug (t=1), and on each subsequent day.

## Acknowledgments

The authors would like to thank their principal investigator Bruce R. Levin for his kibitzing on this project. We would also like to thank Andrew P. Smith and Fernando Baquero for their comments and suggestions on this manuscript.

## Transparency Declarations

The authors have no competing interests to declare.

